# A unique mode of nucleic acid immunity performed by a single multifunctional enzyme

**DOI:** 10.1101/776245

**Authors:** S. M. Nayeemul Bari, Lucy Chou-Zheng, Olivia Howell, Katie Cater, Vidya Sree Dandu, Alexander Thomas, Barbaros Aslan, Asma Hatoum-Aslan

## Abstract

The perpetual arms race between bacteria and their viruses (phages) has given rise to diverse immune systems, including restriction-modification and CRISPR-Cas, which sense and degrade phage-derived nucleic acids. These complex systems rely upon production and maintenance of multiple components to achieve anti-phage defense. However, the prevalence and effectiveness of much simpler, single-component systems that cleave DNA remain unknown. Here, we describe a novel mode of nucleic acid immunity performed by a single enzyme with nuclease and helicase activities, herein referred to as Nhi. This enzyme provides robust protection against diverse staphylococcal phages and prevents phage DNA accumulation in cells stripped of all other known defenses. Our observations support a model in which Nhi acts as both the sensor and effector to degrade phage-specific replication intermediates.

Importantly, Nhi homologs are distributed in diverse bacteria and exhibit functional conservation, highlighting the versatility of such compact weapons as major players in anti-phage defense.

## Introduction

Phages are the most abundant entities in the biosphere (Bergh et al., 1989), and as such they impose a tremendous selective pressure upon their bacterial hosts. Phages attach to a specific host, inject their genetic material, and utilize the host’s enzymes and energy stores to replicate exponentially in a process that typically leads to cell lysis and death. In response to this constant threat, bacteria have evolved an impressive collection of immune systems that undermine nearly every step of the phage infection cycle (Hampton et al., 2020). Such systems may block phage genome entry, interfere with phage DNA replication/expression, and/or as a last resort, precipitate programmed cell death, a process known as abortive infection (Abi), to prevent phages from spreading to neighboring bacteria in the population (Lopatina et al., 2020). Abi can be achieved through a variety of mechanisms and constitutes a remarkably common defense strategy. Indeed, recent years have witnessed a surge of reports on novel bacterial immune systems (Bernheim et al., 2021; Cohen et al., 2019; Doron et al., 2018; Gao et al., 2020; Kronheim et al., 2018; Millman et al., 2020), and many of these ultimately cause cell death. Notable examples utilize RNA-modifying enzymes (Gao et al., 2020), retrons (Gao et al., 2020; Millman et al., 2020), and small molecules (Cohen et al., 2019) as the basis for defense.

As a more direct, and perhaps more effective approach to stemming a phage infection, bacteria employ different systems that sense and destroy phage genetic material. Such systems exhibit a range of complexities, from the simpler restriction-modification (RM) systems to the more sophisticated adaptive immune systems that rely upon clusters of regularly-interspaced short palindromic repeats (CRISPRs) and CRISPR-associated (Cas) genes (Hampton et al., 2020). RM systems are innate immune systems that require at least two components for defense—a nuclease that cleaves specific DNA sequences in the phage, and a DNA methyltransferase that modifies and protects the host genome from cleavage. Further, more complex RM-like systems have recently been described, including Pgl (phage growth limitation) (Hoskisson et al., 2015; Sumby and Smith, 2002), DISARM (defense island associated with restriction-modification) (Ofir et al., 2017), and BREX (bacteriophage exclusion) (Goldfarb et al., 2015; Gordeeva et al., 2019). All of these encode DNA methyltransferases in conjunction with a variety of genes such as proteases, phosphatases, and phospholipases which provide other necessary functionalities for defense. CRISPR-Cas systems are even more elaborate—they retain a molecular memory of phage-derived nucleic acids by capturing small stretches of their genetic material and storing them in the CRISPR locus as “spacers” in between DNA repeats (Hille et al., 2018). The spacer sequences are transcribed to generate small CRISPR RNAs (crRNAs), which are subsequently used in combination with one or more Cas nucleases to identify and eliminate phages harboring sequences complementary to the crRNAs. These systems adapt to new and evolving threats by continually capturing new spacers from new phage invaders. There are two classes, six types and 33 subtypes of CRISPR-Cas systems currently described (Makarova et al., 2020), and different types have been shown to work together (Deng et al., 2013; Hoikkala et al., 2021; Pinilla-Redondo et al., 2020; Silas et al., 2017) and even synergize with RM systems (Dupuis et al., 2013) to ensure a more effective defense. Such added layers have obvious advantages in protecting against diverse and evolving phage predators; however, this strategy comes with the energetic cost of producing and maintaining multiple components, as well as the risk that damage or loss of a single part may render the system inactive. Thus, it is reasonable to speculate that bacteria also employ much simpler, single-component systems to target and degrade phage nucleic acids. However, the prevalence and effectiveness of such minimal systems remain poorly understood.

Here, we describe the discovery and characterization of a unique mode of nucleic acid immunity performed by a single enzyme, herein referred to as Nhi. This enzyme provides full protection against diverse staphylococcal phages, and it is sufficient to prevent phage DNA accumulation in a staphylococcal strain devoid of all other known defenses. In a purified system, Nhi dimerizes and possesses both nuclease and helicase activities. Further, we found that a single-stranded DNA binding protein encoded in one group of phages can protect against Nhi’s effects. Based on these observations, we propose a model in which Nhi senses and degrades phage-specific replication intermediates. Importantly, Nhi homologs can be found in diverse bacterial phyla, and we provide evidence that they are also dedicated to immunity. Our findings highlight the versatility of such compact systems as powerful weapons in anti-phage defense.

## Results

### *SERP2475* protects against diverse staphylococcal phages

We undertook this study with an aim to uncover new mechanisms of immunity in the commensal opportunistic pathogen *Staphylococcus epidermidis* RP62a (Christensen et al., 1987). This organism harbors a Type III-A CRISPR-Cas system (Marraffini and Sontheimer, 2008), an Abi mechanism (Depardieu et al., 2016), and a putative Type I RM system, all of which are encoded within ~30,000 nucleotides of each other (Fig. 1A). Their close proximity is consistent with recent reports that show prokaryotic immune systems typically cluster together within discrete genomic loci known as defense islands (Doron et al., 2018; Gao et al., 2020; Makarova et al., 2011). Importantly, key insights into CRISPR-Cas and Abi in this organism were revealed by studying their molecular interactions with temperate phages ΦNM1 and CNPx, respectively, closely related phages that belong to the family *Siphoviridae* (Depardieu et al., 2016; Goldberg et al., 2014). Indeed, siphophages are the most common members in staphylococcal phage collections (Oliveira et al., 2019), and we reasoned that the identification of new immunity mechanisms might necessarily require the examination of more diverse members. Toward that end, we captured and characterized four new lytic *S. epidermidis* phages belonging to the family *Podoviridae*: Andhra, JBug18, Pontiff, and Pike (Cater et al., 2017; Culbertson et al., 2019). These phages share over 95% sequence identity and the same 20 genes (Fig S1); however, we noticed that they have distinct host ranges—while Andhra and Pontiff can infect the wild-type RP62a strain, JBug18 and Pike can only infect a mutant variant, LM1680 (Jiang et al., 2013), which has a large (~300k nucleotide) deletion encompassing the defense island and neighboring regions (Fig. 1A). These observations led to the hypothesis that JBug18 and Pike are sensitive to genetic element(s) within the defense island.

**Figure 1.**
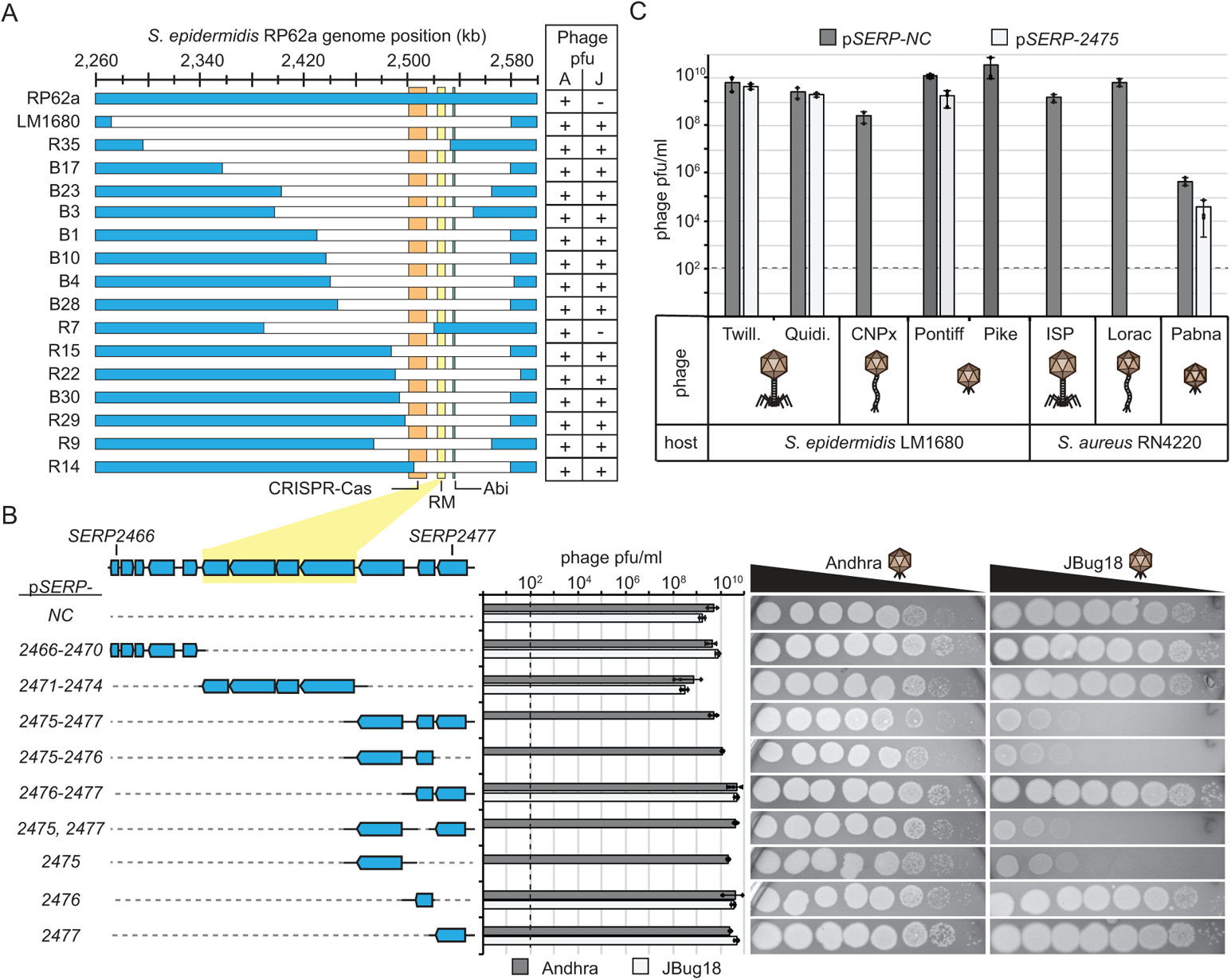
*SERP2475* provides robust immunity against diverse staphylococcal phages. (**A**) A segment of the S. *epidermidis* RP62a genome and deletion mutants. Regions encoding CRISPR-Cas, RM, and Abi systems are highlighted. Strains were challenged with Andhra (A) and JBug18 (J), and resulting plaque-forming units per milliliter (pfu/ml) are indicated: +, ~1 x 10^9^ pfu/ml; -, 0 pfu/ml. (**B**) Magnified view of the genomic region responsible for immunity and corresponding plasmids that were created to encompass them. *S. epidermidis* LM1680 strains harboring indicated plasmids were challenged with phage, and the resulting pfu/ml are shown as an average of triplicate measurements (± S.D.). Images on the right show representative plates following the application of ten-fold dilutions of Andhra and JBug18 (1×10^0^ – 1×10^−7^) atop *S. epidermidis* LM1680 strains bearing indicated plasmids. (**C**) *S. epidermidis* and *S. aureus* strains were challenged with phages representing all three morphological families of tailed phages, and resulting pfu/ml are shown. The dotted line indicates the limit of detection for phage challenge assays. See also Figure S1.

To test this, we used a set of *S. epidermidis* RP62a mutants that bear deletions of varying extents across the defense island (Jiang et al., 2013) (Fig. 1A). These strains were challenged with Andhra and JBug18, representatives from the set of four with resistant and sensitive phenotypes, respectively. The resulting zones of bacterial growth inhibition (plaques) were enumerated, revealing that only one of the deletion mutants (R7) encodes the gene(s) required for complete protection against JBug18. These observations narrowed the protective genetic element(s) to a stretch of ~12,000 nucleotides containing 12 genes (designated as *SERP2466*-*SERP2477*) which incidentally encompasses the RM system (Fig. 1B). To determine which of the 12 gene(s) are responsible for immunity, they were inserted (individually and in groups) into a derivative of plasmid pC194 (Ehrlich, 1977) (herein referred to as p*SERP-*), introduced into *S. epidermidis* LM1680, and the resulting strains were challenged with Andhra and JBug18. Through this analysis, we found that a single gene of unknown function, *SERP2475* (new locus tag *SERP_RS12125*) is sufficient to protect against JBug18 (Fig. 1B). A repeat of this assay with phages Pontiff and Pike showed results similar to those observed with Andhra and JBug18, respectively—while *SERP2475* has no effect on Pontiff, it completely protects against Pike (Fig. 1C).

To further understand the breadth of protection this gene affords, we first challenged the *S. epidermidis* LM1680/p*SERP-2475* strain with additional phages from our collection from different morphological families—*Herelleviridae* (Barylski et al., 2020) (formerly *Myoviridae*) and *Siphoviridae*. We observed that although *SERP2475* has no noticeable impact on the lytic myophages Twillingate and Quidividi (Freeman et al., 2019), it fully protects against siphophage CNPx (Fig. 1C). We also tested the effectiveness of SERP2475 against *S. aureus* phages representing all three families of tailed phages and found that while SERP2475 causes only modest reduction in plaques of podophage Pabna (Culbertson et al., 2019), it affords full protection against myophage ISP (Vandersteegen et al., 2011) and siphophage Lorac (Antoine Marc et al., 2019). Taken together, these data demonstrate that *SERP2475* is sufficient to provide robust protection against diverse staphylococcal phages.

### SERP2475 homologs are distributed in diverse bacteria and exhibit anti-phage activity

We next assessed the distribution of SERP2475 homologs and the extent to which they are functionally conserved. We used tBLASTn to query three NCBI databases (prok_complete_genomes, refseq_genomes, and nt) and identified 302 homologs in distinct genetic backgrounds (Table S1). Although homologs were present in <1% of the fully assembled microbial genomes surveyed, representatives could be found in three bacterial phyla: *Firmicutes*, *Bacteroidetes*, and *Proteobacteria*. Additionally, two of the homologs were found in two *Streptococcus* phages. To better understand their relationships to one another, we generated a phylogenetic tree from 100 selected representatives that encompass the phylogenetic diversity of the group (Fig. 2A and S2, and Supplementary File 1). This analysis revealed that the homologs cluster into three distinct clades that are somewhat incongruent with host phylogeny--While clade III contains homologs strictly found in *Proteobacteria*, clades I and II contain homologs originating from two of the three phyla. This observation suggests that *SERP2475* and its homologs are likely disseminated through horizontal gene transfer. Supporting this, many of the homologs are encoded on plasmids (Fig. 2A and Table S1). To determine their level of functional conservation, we first checked for proximity to genes with known defense functions. Amino acid sequences of the thirty proteins encoded upstream and downstream of each homolog were extracted and searched for identifiable protein domains using the hmmer software v.3.3.2 (hmmer.org). The predicted protein families (pfams) of these flanking proteins were then searched against 306 pfams with known functions in anti-phage defense (Table S2). Figure 2B shows the fraction of homologs that have between one and ten defense-related neighbors encoded within expanding windows of 10, 20, and 30 genes. This data revealed that a significant fraction of SERP2475 homologs are indeed encoded proximal to other known defenses. For instance, 72% of homologs have at least one defense neighbor encoded within 20 genes (Fig. 2B and Table S3), thus supporting roles in anti-phage defense. The most frequently encountered defense neighbors within 20 genes include proteins with TOPRIM (PF01751), ATPase (PF13304, PF00176, and PF00004) and methyltransferase (PF02384 and PF01420) domains (Fig. 2C and Table S4).

**Figure 2.**
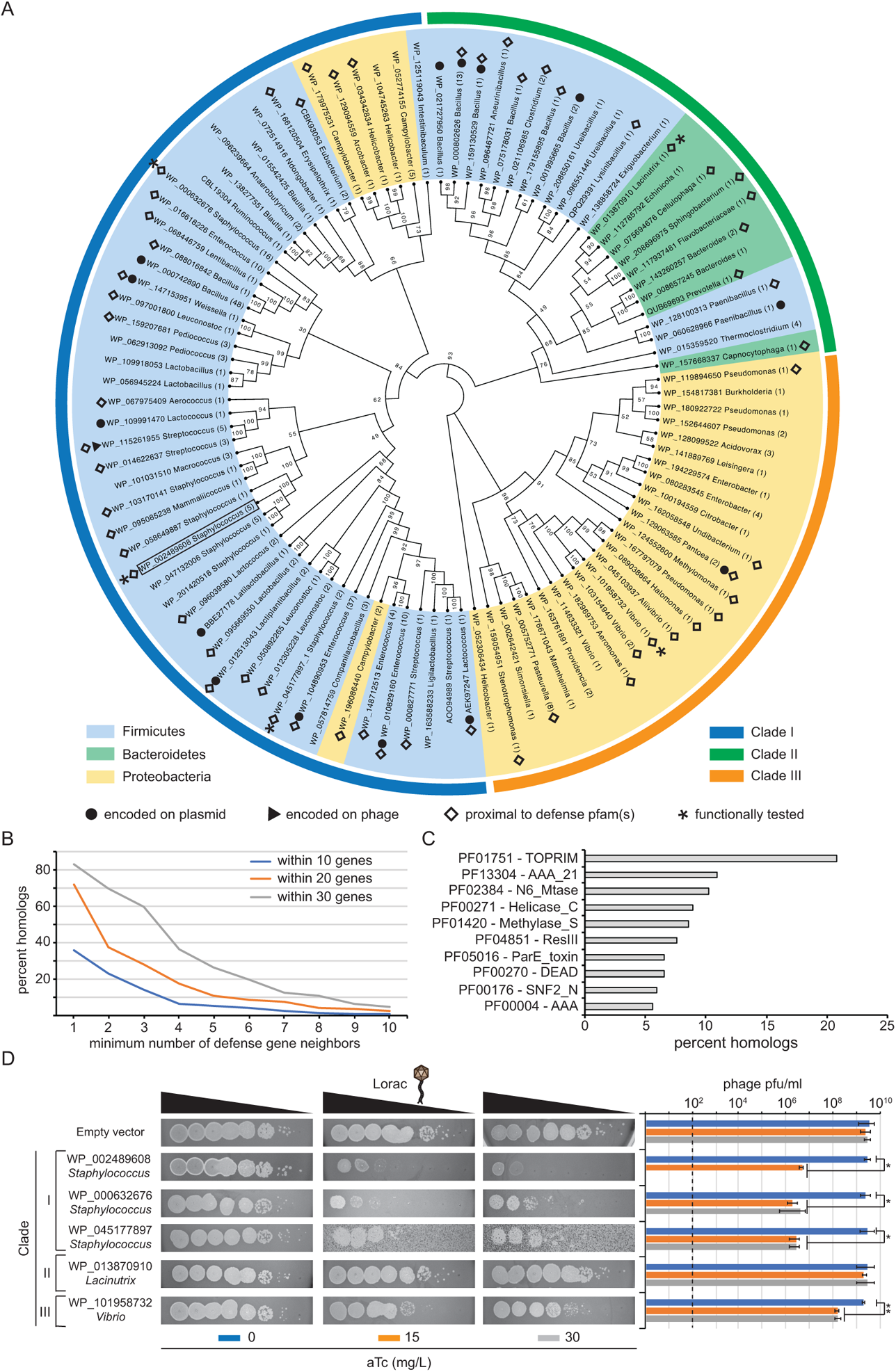
SERP2475 homologs exhibit functional conservation. (**A**) A cladogram showing the phylogenetic distribution of SERP2475 and selected homologs. Branch labels indicate bootstrap values and tip labels show the NCBI RefSeq ID number, genus from which the homolog originated, and number of distinct species/strains within the same genus that harbor an identical or closely-related homolog (>90% amino acid identity). Inner ring highlighting indicates the phylum, and the outer ring coloration indicates the clade (I-III) in which each homolog belongs. Representatives encoded on plasmids (closed circles), phages (closed triangle), and within 20 genes of at least one neighbor with a predicted defense function (open diamonds) are indicated. Asterisks mark homologs that are functionally characterized in this study, and SERP2475 is additionally enclosed with a box. (**B**) A neighborhood analysis showing the fraction of homologs (including SERP2475, n=303) that have between 1 and 10 neighbors dedicated to anti-phage defense within 10, 20, and 30 flanking genes. (**C**) Top ten defense-related protein families (pfams) encoded within 20 genes of SERP2475 homologs. (**D**) Functional analysis in which the coding sequences of selected homologs were inserted into a plasmid downstream of an anhydrotetracycline (aTc) inducible promoter and subsequently introduced into *S. aureus* RN4220. Images show ten-fold dilutions of phage Lorac (10^0^ – 10^−7^) spotted atop lawns of indicated strains grown in the absence or presence of aTc (15 and 30 mg/L). An average of triplicate measurements of pfu/ml (± S.D) are shown as a representative of three independent trials. Asterisks indicate p-values <0.05 (*) and <0.0005 (**) in a two-tailed t-test. The dotted line shows the limit of detection for this assay. See also Figure S2.

To further explore the functional conservation of these homologs, we selected four representatives from the three clades and tested for anti-phage activity (Figs. 2A and S2, and Table S1). Importantly, these homologs share minimal (32-36%) amino acid sequence similarity when compared with SERP2475. The coding sequences for SERP2475 and selected homologs were inserted into a plasmid (herein referred to as pTET-) downstream of an anhydrotetracycline (aTc) inducible promoter and then introduced into *S. aureus* RN4220. The resulting strains were challenged with *Siphoviridae* phage Lorac in the presence and absence of the inducer. As expected, SERP2475 (RefSeq ID WP_002489608) affords full protection against the phage when the cells were grown in the presence of the maximum concentration of aTc (Fig. 2D). Although expression of the Clade II homolog (WP_013870910 from phylum *Bacteroidetes*) resulted in no detectable decrease in plaque size or number, the two *Staphylococcus* homologs from Clade I (WP_000632676 and WP_045177897) and the *Vibrio* homolog from Clade III (WP_101958732 from phylum *Proteobacteria*) caused significant reduction in plaque numbers (1000- to 10-fold, respectively). A noticeable decrease in plaque size was also observed in the latter. Interestingly, one of the staphylococcal homologs (WP_045177897) also appeared to be toxic to the cells, as evidenced by the ‘unhealthy’ appearance of the lawn when cells harboring the homolog are grown in the presence of inducer (Fig. 2D). Altogether, these data provide evidence that distant homologs of SERP2475 are also dedicated to anti-phage defense.

### *SERP2475* limits phage DNA accumulation

We next sought to investigate the mechanism of immunity, and began by assessing which stage of the phage infection cycle SERP2475 targets. As previously mentioned, common strategies that bacteria employ in anti-phage defense include masking/modifying the cell surface to prevent phage attachment, targeting and degrading phage-derived nucleic acids, and causing programmed cell death through various Abi mechanisms (Fig. 3A). To test if *SERP2475* hinders phage attachment, we conducted an adsorption assay in which *S. epidermidis* LM1680 cells bearing p*SERP-2475* were combined with a defined number of phage particles for ten minutes— just enough time for phages to attach to cells, but not long enough for these phages to complete their replication cycle (Cater et al., 2017). The mixture was then pelleted to remove both the cells and phages attached to them, and the phages remaining in suspension were enumerated to determine the number that adsorbed (i.e. attached) to cells. This assay revealed that JBug18 attaches to LM1680/p*SERP-2475* just as efficiently as Andhra (Fig. 3B), thus ruling out an adsorption-blocking mechanism.

**Figure 3.**
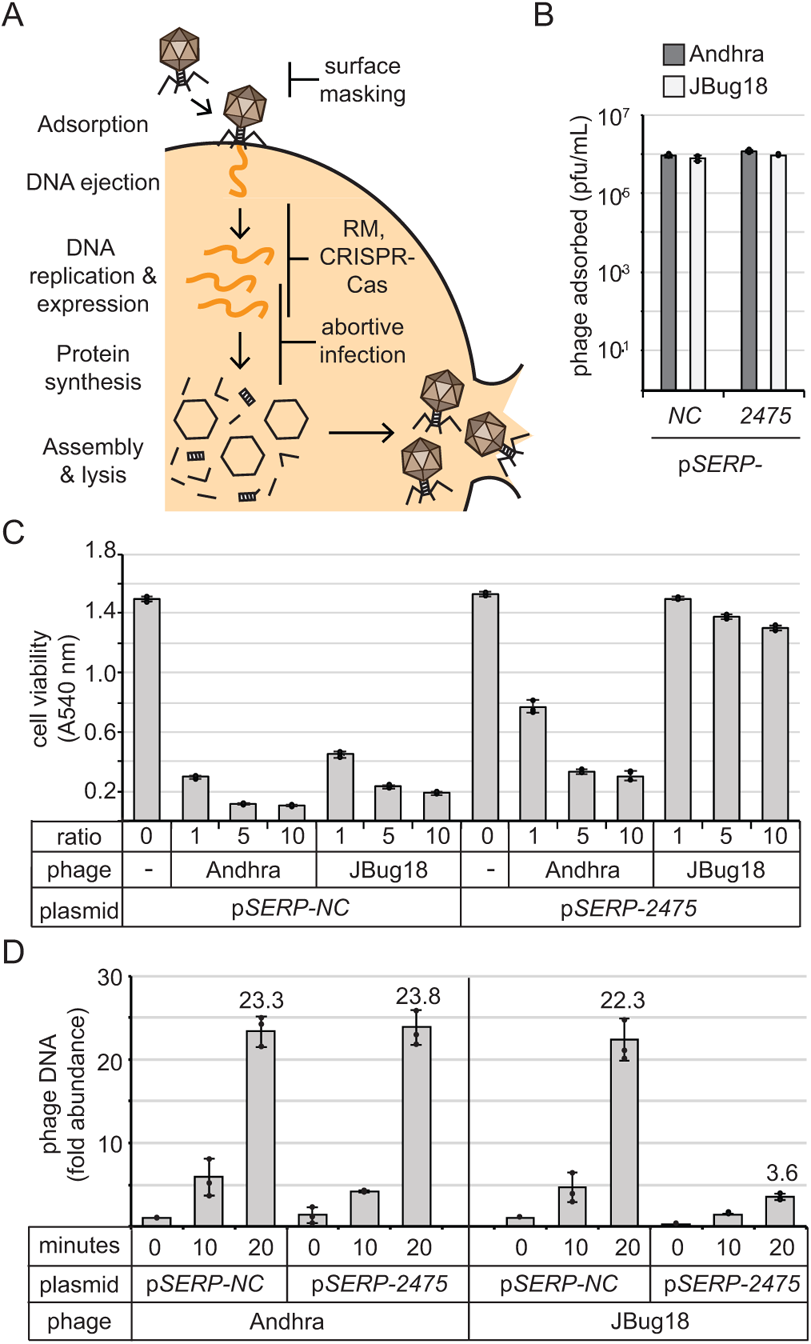
SERP2475 impairs phage DNA accumulation. (**A**) Illustration of the lytic phage replication cycle and common defense systems that interfere with each step. (**B** and **C**) Results of an adsorption assay (**B**) and a cell viability assay (**C**) following challenge of *S. epidermidis* LM1680 cells bearing indicated plasmids with Andhra and JBug18. (**D**) Relative abundance of phage DNA at various time points following phage infection as measured by qPCR. For all experiments, the mean ± S.D. of triplicate measurements are shown as a representative of at least two independent trials.

A cell viability assay was next performed to test for Abi. The prediction is that if programmed cell death accompanies immunity, then challenge with a high proportion of phages to cells (≥1:1) would lead to significant decline in cell viability similar to that observed in the absence of immune protection (Goldfarb et al., 2015; Ofir et al., 2017). To test this, phages were combined with LM1680/p*SERP-2475* cells in liquid media at ratios of 1:1, 5:1, or 10:1, and cell viability was measured following 5 hours of incubation. We found that while Andhra causes significant death of LM1680/p*SERP-2475*, JBug18 elicits only a minor decrease in the viability of this strain, even when phages outnumber bacteria 10:1 (Fig. 3C). These observations suggest that cell suicide is unlikely to be the mechanism by which *SERP2475* affords protection.

Finally, we tested whether *SERP2475* impacts phage DNA levels in the cell. Quantitative PCR (qPCR) was used to track the accumulation of phage DNA at various time points following infection with both phages. The results showed that while Andhra’s DNA accumulates to ~20-fold by 20 minutes post-adsorption in LM1680/p*SERP-2475*, JBug18’s DNA accumulates to less than 4-fold in the same time period (Fig. 3D). These observations support a hypothesis whereby SERP2475 protects against phages by directly interfering with phage DNA replication and/or expression.

### SERP2475 relies upon nuclease and helicase activities to perform immunity

To begin to understand the catalytic function of this protein, we first conducted *in silico* analyses. *SERP2475* encodes a 606 amino acid protein, and according to Interproscan (Blum et al., 2021), a web-based tool that predicts protein domains and other important functional sites, the most prominent features of SERP2475 are a central domain of unknown function (DUF) 2075 (protein family PF09848) and a P-loop NTPase domain that overlaps with it (Fig. 4A). Additionally, the structural homology search tools HHPred (Zimmermann et al., 2018) and Phyre2 (Kelley et al., 2015) identified a handful of superfamily 1 helicases as close homologs, including T4 phage Dda helicase (He et al., 2012) (E-value 1.2 x 10^−21^). The latter tools also predicted structural similarities between the N-terminus of SERP2475 and a putative HsdR restriction endonuclease from *Vibrio vulnificus* (Uyen et al., 2009) (E-value 1.1 x 10^−6^). Although SERP2475 shares minimal amino acid sequence similarity in pairwise comparisons with full length HsdR_Vv and Dda (19% and 22%, respectively), we located conserved residues in key motifs corresponding to a putative PD-(D/E)XK nuclease domain on its N-terminus (Fig. 4B) and putative helicase domain spanning the central portion of the protein (Fig. 4C). Further, we observed a striking conservation of these critical residues in the sequence alignment with SERP2475 and the 99 representative homologs (Fig. 4D), suggesting that these domains are likely important for protein function. To confirm, we constructed two mutant versions of p*SERP-2475* that encode alanine substitutions in the putative nuclease and helicase domains (asterisks in Fig. 4B and C). The strains were then challenged with phages Andhra and JBug18, and the results showed a complete restoration of JBug18 replication in both mutant strains (Fig. 3E). These collective observations support the hypothesis that SERP2475 and its homologs use nuclease and helicase activities to achieve immunity.

**Figure 4.**
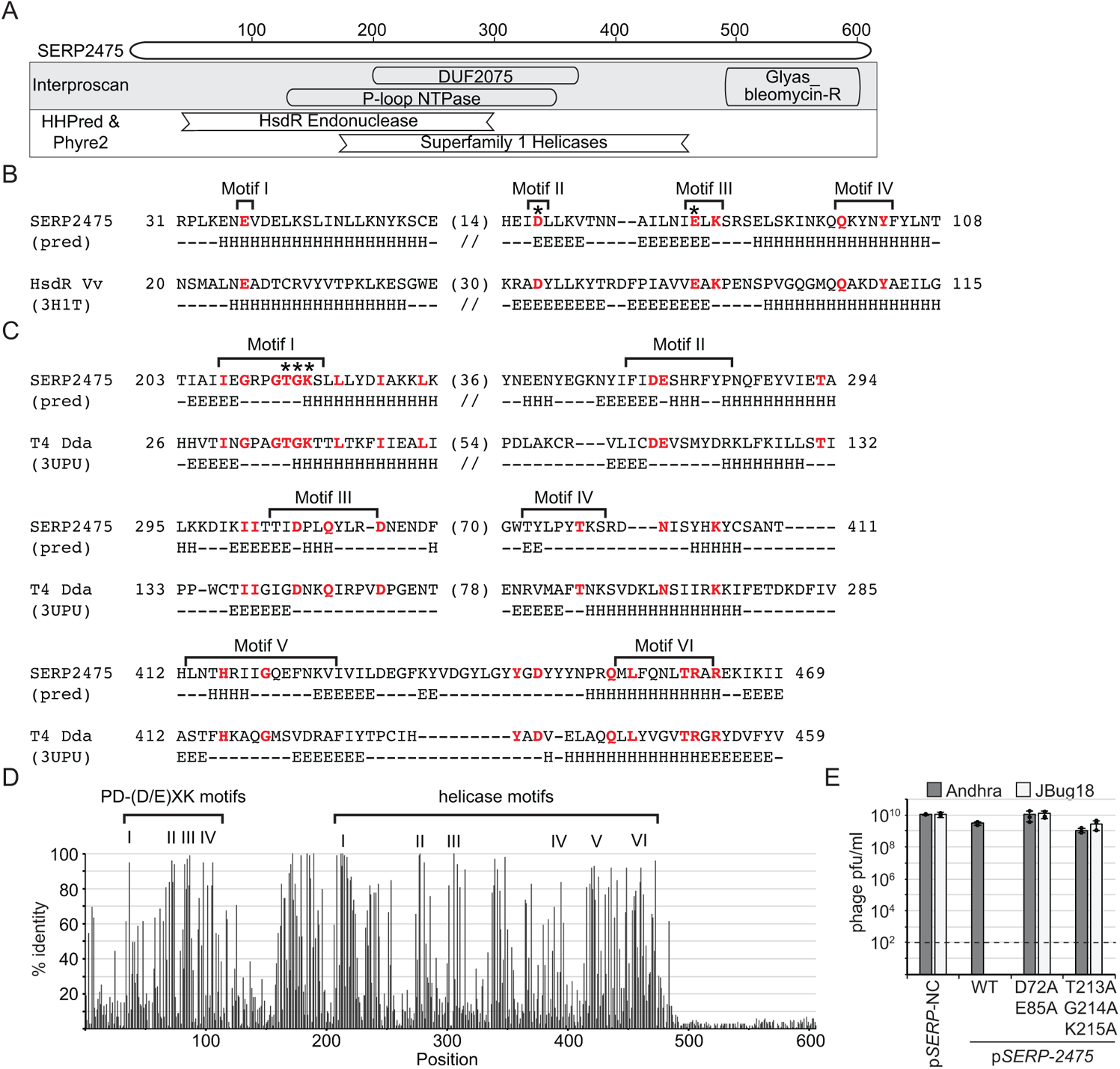
Conserved nuclease and helicase domains are required for immunity *in vivo*. (**A**) Predicted domains and structural homologs of SERP2475. (**B** and **C**) Pairwise sequence alignments between SERP2475 and relevant regions of the putative HsdR endonuclease from *Vibrio vulnificus* and the Dda helicase from phage T4 (Swiss PDB IDs 3H1T and 3UPU, respectively) as determined by HHPred. Residues colored red indicate positions of sequence identity and asterisks mark residues that were subjected to mutational analysis in this study. Predicted (pred) and actual alpha helices (H) and beta sheets (E) are indicated below each position in the alignments. (**D**) The fraction of 99 homologs that possess amino acids identical to those of SERP2475 at each position in their multiple sequence alignment. Positions of critical amino acids comprising the PD-(D/E)XK nuclease and helicase motifs are indicated at the top. (**E**) *S. epidermidis* LM1680 strains bearing the indicated plasmids were challenged with phages Andhra and JBug18, and resulting pfu/ml were enumerated. Shown are an average of triplicate measurements (±S.D.). The dotted line indicates the limit of detection for this assay.

To test for these enzymatic functions, we next took a biochemical approach. *SERP2475* was introduced in the pET28b expression vector downstream of an N-terminal 10His-Smt3 tag, overexpressed in *E. coli*, and subjected to a three-step purification process involving two steps of affinity purification followed by size exclusion chromatography (Fig. 5A). During the first two steps, we noticed that SERP2475 co-purifies with a smaller species, and following digestion of the 10His-Smt3 tag by the SUMO protease, both exhibit a reduction in size of ~14 kDa (Fig. 5B). This suggested that the smaller species is also tagged, and thus comprises an N-terminal fragment of the full-length protein. To confirm, we excised the bands in the SDS-PAGE gel corresponding to the full length (~72 kDa) and putative truncated variant (~23 kDa) in the flow-through fraction from the second purification step and subjected the proteins to mass spectrometry analysis. The results indicated that SERP2475 is indeed the most abundant protein in both bands (Tables S5 and S6), with the truncated version showing dense peptide coverage over only the first ~230 amino acids (AAs) (Fig. 5C). The subsequent size exclusion chromatography step successfully separated the full-length and truncated versions, and also revealed that the full-length species forms a dimer in solution, as evidenced by the presence of two adjacent peaks in the chromatogram that contain the pure full-length protein (Fig. 5D).

**Figure 5.**
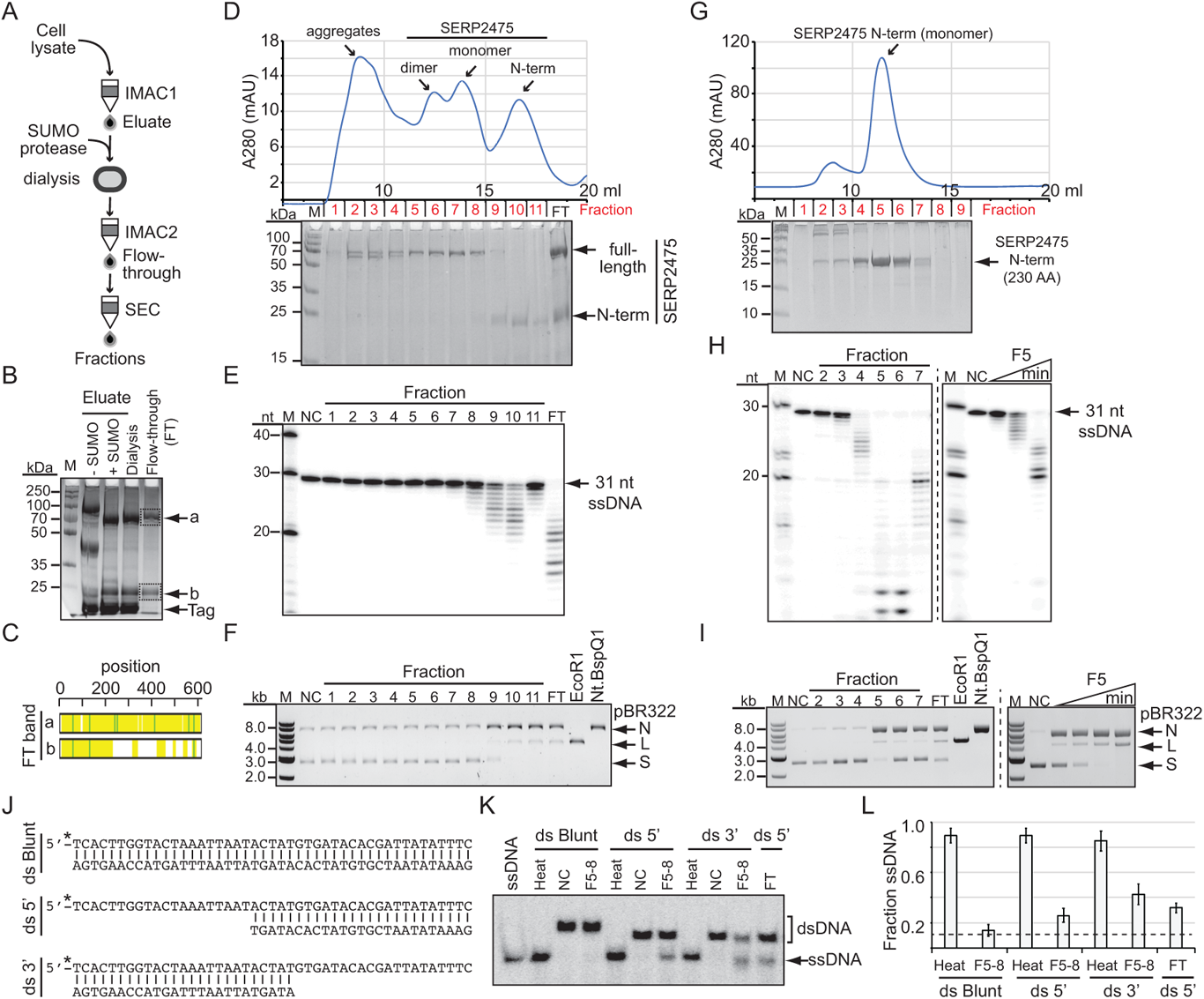
SERP2475 has nuclease and helicase activities. (**A**) Three-step protein purification process used in this study. IMAC, immobilized metal affinity chromatography; SEC, size exclusion chromatography. (**B**) Image of representative SDS-PAGE gel showing protein species present throughout the first two chromatography steps. Dashed boxes encompass protein bands that were excised and subjected to mass spectrometry analysis. (**C**) Peptide coverage of SERP2475 in excised bands. Yellow, identified peptides; green, cysteine carbamidomethylation or methionine oxidation. (**D** and **G**) Size exclusion chromatograms (top) and SDS-PAGE gels (bottom) resolving fractionated proteins in full-length (D) or the N-terminal 230 AAs (G) of SERP2475. mAU, milli absorbance units at 280 nm. (**E** and **F**) Exonuclease (E) and nickase (F) assays in which indicated substrates were combined with fractions from the full-length protein prep in panel D. (**H** and **I**) Left, Exonuclease (H) and nickase (I) assays in which indicated substrates were combined with fractions from the N-terminal 230 AA prep in panel G. Right, time course assays in which substrates were incubated with fraction 5 (F5) in a 13 nM reaction for various amounts of time (5, 10, and 20 min (H) or 30, 60, 120, and 240 min (I)). Products of exonuclease and nickase reactions were resolved on denaturing urea-PAGE and native agarose gels, respectively. For nickase assays, EcoRI and Nt.BspQI were used as controls to generate linear (L) and nicked (N) products, respectively, from the supercoiled (S) plasmid. Shown are representatives of at least three independent trials. (**J**) Substrates used in helicase assays. (**K**) Helicase assays in which indicated substrates were incubated at 37°C for 1 h with or without full-length SERP2475 pooled fractions 5-8 (F5-8) in a 4 nM reaction and resolved using native PAGE. As a positive control for complete unwinding, substrates were heated 95°C for 2 min. (**L**) Quantification of unwound (ssDNA) substrate following heating or incubation with full-length SERP2475 F5-8. Shown is an average of four independent trials (± S.D.). M, molecular weight ladder; kb, kilobase; nt, nucleotide; kDa, kilodalton; NC, negative control, ss, single-stranded; ds, double-stranded. See also Figure S3.

We next conducted *in vitro* functional analyses. Nuclease assays were first performed with the fractionated protein and revealed both 3’-5’ exonuclease and plasmid nicking activities (Fig. 5E and F, respectively) stemming from fractions containing the N-terminal fragment. Interestingly, these activities were nearly absent in fractions containing the full-length protein, suggesting a mechanism of autoinhibition. To confirm that the N-terminal domain is sufficient to produce both activities, we introduced the region of *SERP2475* encoding the first 230 AAs of the protein into the pET28b expression vector downstream of the 10His-Smt3 tag and purified this truncated version to homogeneity using the same purification protocol (Fig. 5A). We observed that the N-terminal fragment elutes as a monomer (Fig. 5G), indicating that the dimerization domain is likely located elsewhere on the protein. Importantly, nuclease assays confirmed that the N-terminal domain is sufficient to produce robust exonuclease and nickase activities (Fig. 5H and I, respectively). The enzyme is unable to further degrade the nicked or linearized double-stranded DNA products when given up to 4 hours of incubation (Fig. 5I), suggesting that supercoiling is essential for cleaving double-stranded DNA. Both activities require either Mg^2+^ or Mn^2+^ and can be observed using different DNA substrates (Fig. S3), providing evidence that these activities are sequence non-specific.

We also tested for helicase activity. As mentioned earlier, the central portion of SERP2475 has conserved helicase motifs and many of its predicted structural homologs are superfamily 1 helicases (Fig. 4). These enzymes are known to function as a monomer or dimer to unwind double-stranded substrates using energy supplied by ATP (Fairman-Williams et al., 2010). To test for this activity, the full-length fractions containing the monomer and dimer peaks (Fig. 5D) were pooled, concentrated, and incubated with ATP, Mg^2+^, and various DNA substrates (Fig. 5J). The results showed that SERP2475 can indeed unwind double-stranded DNA when offered a 5’- or 3’-single-stranded overhang (Fig. 5K and L). Altogether, these results demonstrate that SERP2475 possesses exonuclease and nickase activities stemming from its N-terminus, as well as bidirectional helicase activity.

### A single-stranded DNA binding protein protects against SERP2475 immunity

To further refine our understanding of SERP2475’s targeting specificity, we sought to determine how Andhra resists immunity. Andhra and JBug18 encode the same 20 proteins (Fig. S4A), and a pairwise alignment of their coding regions show that they differ at only 705 positions by either a single-nucleotide polymorphism (SNP) or a gap (Supplementary file 2). To narrow down which SNPs and/or gaps in Andhra are important for resistance to immunity, we first attempted to isolate naturally-evolved JBug18 mutants that can escape immunity by plating concentrated phage preparations with LM1680/p*SERP-2475*. After several failed attempts at recovering plaques, one attempt yielded resistant phages, which upon further inspection were found to possess hybrid genomes that contain a patchwork of Andhra and JBug18 sequences. These hybrids necessarily arose through the inadvertent mixing of the two phages and propagation on the same host strain. Nonetheless, this fortuitous accident proved invaluable in helping to pinpoint the region required for immune resistance—since all hybrids can escape immunity, we reasoned that they must share Andhra-derived sequences in the region(s) required for resistance. To test this, we purified and sequenced eight such hybrids, and determined the fraction of hybrids that possess Andhra identity at each of the 705 differing positions across their coding regions. We found that all eight hybrids share Andhra identity at positions 891-2117 in the alignment (Fig. 6A and Supplementary file 2). This region overlaps gene products (*gp*) *03-06* in the phage genomes and encompasses 69 SNPs and gaps, of which, 64 are concentrated within *gp03* and *gp04* (Fig. 6B). Accordingly, we speculated that one or both of the latter genes are responsible for resistance.

**Figure 6.**
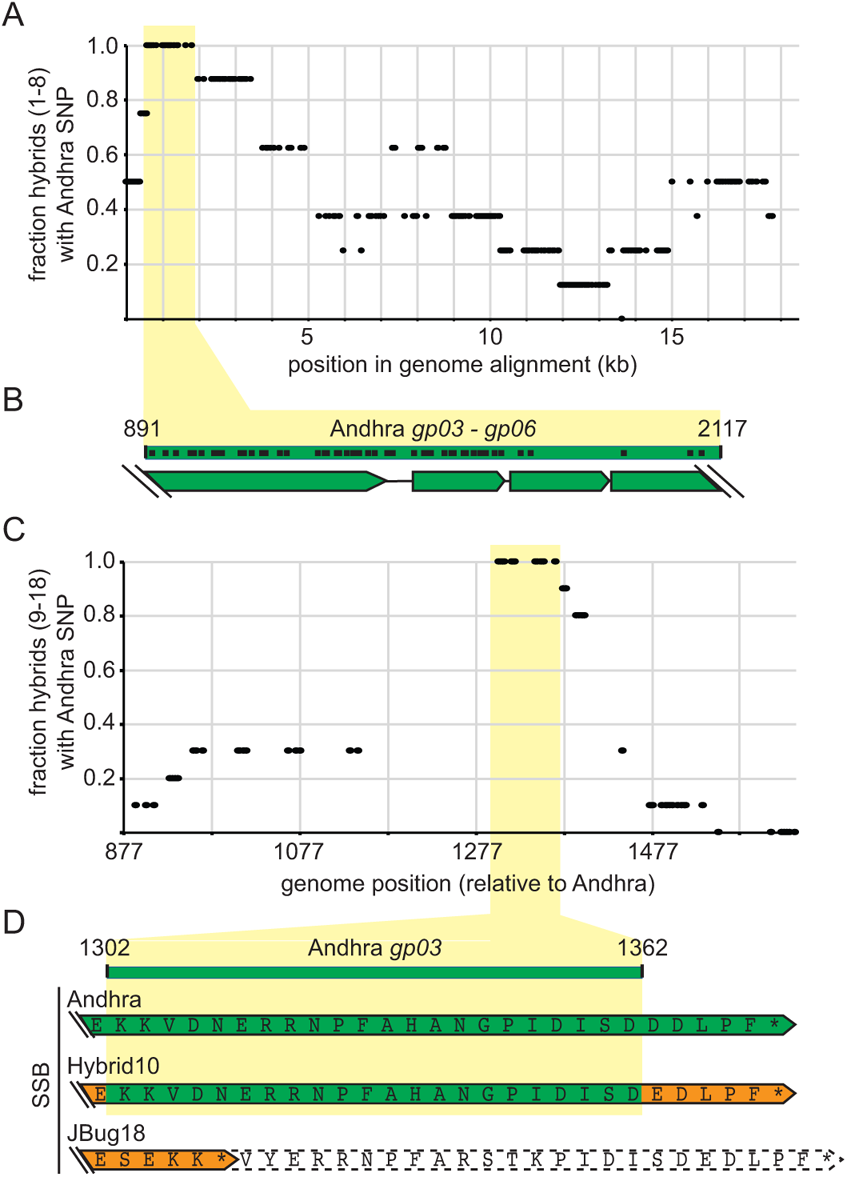
A phage-encoded single-stranded DNA binding protein protects against immune function. (**A** and **C**) Fractions of JBug18-Andhra Hybrids that harbor Andhra identity at positions where the parental phage genomes differ. Segments common to all hybrids are highlighted in yellow. (**B** and **D**) Expanded views of the highlighted common regions. Black dots in (B) indicate relative positions where SNPs and/or gaps occur in an alignment between the parental phages. In (D), green and orange arrows with solid borders delimit phage-encoded SSBs, and the white arrow with dotted border delimits a truncated portion. See also Figure S4.

To narrow down the protective region even further, a second set of resistant JBug18 hybrids were generated which bear Andhra-derived sequences in *gp03* and *gp04*. This was accomplished by introducing Andhra’s *gp03* and/or *gp04* coding regions into *S. epidermidis* LM1680 on plasmids, and then propagating JBug18 on these strains to allow the phage to recombine with the Andhra-derived sequences (Fig. S4B). The resulting phages were then plated on LM1680/p*SERP-2475* to select for resistant phage recombinants. Ten such hybrids (9-18) were purified, and sequencing across *gp03* and *gp04* revealed that they had all acquired a 60-nucleotide stretch spanning positions 1302-1362 in Andhra’s genome (Fig. 6C and Supplementary file 2). This region overlaps *gp03*, which encodes a single-stranded DNA binding protein (SSB, Fig 6D, and Supplementary file 3). Importantly, JBug18 has a 5-nucleotide insertion in this region and consequently harbors a truncated variant of the SSB (Fig. 6D and S4C). However, by acquiring the 60-nucleotide stretch from Andhra, all ten hybrids had restored the reading frame and hence encode a full-length SSB, suggesting that the C-terminus of the SSB is essential for escape from immunity. In agreement with these observations, phages Pontiff and Pike possess the expected *gp03* genotypes: While the resistant phage Pontiff encodes a full-length SSB, the sensitive phage Pike encodes a truncated version, this time due to a single nucleotide deletion (Fig. S4D).

## Discussion

Here, we describe a new mode of nucleic acid immunity performed by a single enzyme with nuclease and helicase activities. Taking into account these demonstrated functions, we propose to name this enzyme and its homologs Nhi (<u>N</u>uclease-<u>h</u>elicase mediated immunity). One attribute that sets Nhi apart from other DNA-targeting immune systems is that it does not rely upon recognition of specific DNA sequences (Fig. 4), and yet abrogates phage DNA accumulation without causing appreciable cell death (Fig. 3). These observations beg the question: What is the basis for phage specificity? Our finding that the SSB in one group of phages can protect against immunity (Fig. 6) supports a preliminary model in which Nhi senses and degrades replication intermediates that are unique to these phages (Fig. 7). Staphylococcal *Podoviridae* phages use a protein-priming mechanism of replication (Salas et al., 2016; Vybiral et al., 2003), in which terminal proteins (TPs) covalently linked to the 5’-ends of their linear dsDNA provide a hydroxyl group upon which to initiate replication (Fig. 7A). The initial stages of replication involve local unwinding of DNA ends by double-stranded DNA binding proteins and subsequent release of the free 3’-end. Under normal circumstances, the 3’-end is captured by the phage DNA polymerase (DNAP) in complex with a second TP, whereupon replication ensues. We speculate that Nhi may compete with DNAP for the free 3’-end and use its 3’-exonuclease and helicase activities to processively degrade the phage genome. SSBs are known to bind and protect DNA and regulate the replication machinery, particularly through interactions with their C-terminus (Shereda et al., 2008). The precise mechanism by which the C-terminus of the *Podoviridae* phages’ SSB protects against Nhi’s effects remains to be determined.

**Figure 7.**
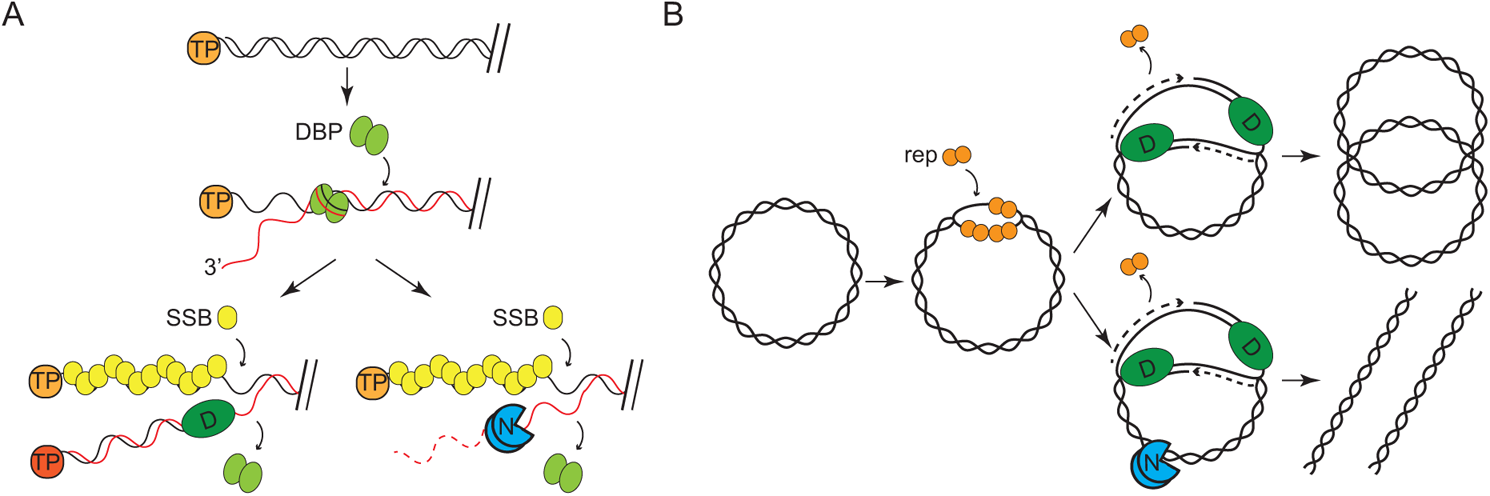
Proposed mechanism for SERP2475 (Nhi) immunity. (**A**) Protein-priming mechanism of replication in staphylococcal *Podoviridae* phages. (**B**) Theta replication in staphylococcal *Siphoviridae* phages. TP, terminal protein; DBP, double-stranded DNA binding protein; rep, replication initiator protein; D, DNA polymerase; N, Nhi. Refer to the discussion for more details.

Remarkably, phages with very different mechanisms of replication also succumb to Nhi. These include the Twort-like phage ISP, and lambda-like phages Lorac and CNPx (Fig. 1). Although the replication mechanism of the former remains poorly understood (Klumpp et al., 2010), the latter undergo several rounds of theta replication early in the infection cycle, followed by rolling-circle replication (Casjens and Hendrix, 2015; Narajczyk et al., 2007) (Fig. 7B). We speculate that Nhi may also recognize and nick supercoiled theta replication intermediates. Since the full-length Nhi forms a dimer in solution (Fig. 5D), it has the potential to introduce two adjacent nicks on opposing strands and cause double-stranded breaks in the phage genome. Indeed, even the monomeric 230 AA fragment caused some plasmid linearization (Fig. 5F and I), presumably through simultaneous nicking on opposite strands that co-localized by chance. Since full-length Nhi is largely devoid of nuclease activity *in vitro* (Fig. 4), we speculate such phage-specific replication intermediates, perhaps in combination with one or more phage-encoded proteins bound to them could constitute recognition elements that trigger Nhi activation.

It is unclear how the full length version of Nhi is truncated within the cells prior to purification (Fig. 5B and C), and we cannot exclude the compelling possibility that generation of the truncated form might constitute a natural part of its mechanism of action. Nonetheless, its unexpected appearance allowed us to reconstitute the nuclease activities stemming from Nhi’s N-terminal PD-(D/E)XK domain. These domains are found in a broad superfamily of metal-dependent nucleases involved in diverse functions, including DNA restriction, recombination, and repair (Steczkiewicz et al., 2012). Members of this group include restriction enzymes, holliday junction resolvases, herpesvirus exonucleases, and many others from in all kingdoms of life. Nhi’s helicase motifs comprise a separate domain that overlap with the DUF2075 (Fig. 4). In addition to the phage T4 Dda helicase, close predicted structural homologs include the human helicase Upf1 (E-value 2.5 x 10^−23^), and mouse nuclease-helicase DNA2 (E-value 1.2 x 10^−21^), which are involved in nonsense-mediated mRNA decay and DNA replication/repair, respectively (Chakrabarti et al., 2011; Zhou et al., 2015). All three are superfamily 1B helicases, which unwind double-stranded substrates in the 5’-3’ direction. Interestingly, Nhi’s helicase activity is bipolar (Fig. 5K), which undoubtedly allows it to act on more diverse substrates. We also noted the presence of a C-terminal domain (Glyas_Bleomycin-R) that is poorly conserved across Nhi homologs (Fig. 4A and D). Such domains are found in a group of metalloenzymes that perform a variety of activities, including isomerizations, epimerizations, and oxidative cleavage of C-C bonds (Armstrong, 2000). Whether and how this domain coordinates with the others to achieve immunity are subjects of ongoing work.

In contrast to the *S. epidermidis* Nhi, other homologs tested showed more modest anti-phage activity (Fig. 2). This could possibly be explained by incompatibility with the foreign phage target or heterologous host background, which together might lessen their apparent effectiveness. In support of this, the homologs that were most effective against *S. aureus* RN4220 phage Lorac originated from two different *S. aureus* strains. Nonetheless, the observation that the *V. vulnificus* homolog could still afford protection is remarkable and underscores the versatility of this mode of prokaryotic immunity. Interestingly, one of the *S. aureus* homologs abrogated cell growth while also blocking phage replication (Fig. 2), suggesting that it may cause collateral cleavage of host DNA, and as a consequence, slow cell division. Indeed, the latter represents a valid strategy for nucleic acid immunity, as some CRISPR-Cas systems are known to simultaneously damage phage and host nucleic acids and cause cell dormancy and/or death during persistent phage infection (Meeske et al., 2019; Ofir et al., 2017; Rostøl and Marraffini, 2019; Watson et al., 2019).

Finally, it bears mentioning that the SERP2475 homologs identified in this study represent but a small subset of DUF2075 domain-containing proteins. According to Interproscan, these constitute a large family of conserved proteins with over 7,000 members that can be found in organisms spanning all domains of life. Although the majority of members are encoded in bacteria, many are found in eukaryotes, a handful of which are in humans. Of these, the Schlafen (slfn) family proteins Slfn5, Slfn11, and Slfn13 have been shown to restrict the replication of diverse viruses, including human immunodeficiency virus (HIV) and herpes simplex virus (HSV) (Kim et al., 2021; Li et al., 2012; Valdez et al., 2019; Yang et al., 2018). Notably, Slfn13 also relies upon endonuclease activity for anti-viral defense (Yang et al., 2018). Such functional conservation across phylogenetic boundaries has become a recurring theme in recently-described defense systems (Bernheim et al., 2021; Cohen et al., 2019; Kazlauskiene et al., 2017; Niewoehner et al., 2017), and we anticipate continued investigation of prokaryotic DUF2075 proteins has the potential to seed new insights into human immunity.

## Methods

### Bacterial strains and growth conditions

S. *epidermidis* RP62a and mutant variants were a generous gift from Luciano Marraffini. *S. epidermidis* strains were grown in Brain Heart Infusion (BHI, BD Diagnostics), *S. aureus* strains were grown in Tryptic Soy Broth (TSB, BD Diagnostics), *E. coli* DH5α was grown in Luria Bertani (LB) broth (VWR), and *E. coli* Rossetta2 (DE3) was grown in Terrific Broth (TB, VWR) for protein purification. Growth media was supplemented with the following: 10 µg/ml chloramphenicol (to select for pC194-based plasmids), 10 µg/ml tetracycline (to select pT181-based plasmids), 10 µg/ml erythromycin (to select for pTET-based plasmids), 15 µg/ml neomycin (to select for *S. epidermidis* cells), 30 µg/ml chloramphenicol (to select for *E. coli* Rossetta2 plasmids) and 50 µg/ml kanamycin (to select for pET28b-10HisSmt3-based plasmids).

### Phage propagation and plate infection assays

*S. epidermidis* phages (Andhra, JBug18, Pontiff, Pike, Quidividi, and Twillingate) and *S. aureus* phages (ISP, Lorac, and Pabna) were propagated on their respective hosts to create stocks. Stock concentrations in plaque-forming units per ml (pfu/ml) were determined as previously described (Antoine Marc et al., 2019; Cater et al., 2017; Culbertson et al., 2019; Freeman et al., 2019). For plate infection assays, 10-fold dilutions of phage stocks were spotted atop a lawn of indicated strains using the double-agar overlay method (Cater et al., 2017). For assays using the anhydrotetracycline-(aTc-) inducible system (Fig. 2), plates and top agar were supplemented with 15 or 30 mg/L aTc, as indicated. Following overnight incubation, phage pfu/ml were enumerated.

### Construction of pC194, pT181, and pTET-based plasmids and introduction into staphylococcal strains

All pC194-, pT181-, and pTET*-*based plasmids were constructed using either inverse PCR or Gibson Assembly(Gibson et al., 2009) with the primers listed in Table S7 as described previously (Walker et al., 2017). pAH011 (Hatoum-Aslan et al., 2011), a derivative of pC194 (Ehrlich, 1977), was used as the backbone for plasmids designated as p*SERP-*in this study. Plasmid pT181 (Khan et al., 1981) was used as backbone for pT181*-gp03* and pT181*-gp0304*. pTarget (Samai et al., 2015) was used as the backbone for all plasmids designated as pTET-in this study. All assembled plasmids were first introduced into *S. aureus* RN4220 (pC194- and pTET*-*based plasmids) or OS2 (pT181-based plasmids) via electroporation, and inserted sequences were confirmed by PCR amplification and Sanger sequencing (performed by Eurofins MWG Operon) using primers shown in Table S7. Confirmed plasmids were purified using EZNA Plasmid Mini Kit (Omega Bio-tek), and where indicated, introduced into *S. epidermidis* LM1680 via electroporation as previously described (Walker et al., 2017).

### Homolog identification

To identify homologs, the amino acid sequence of SERP2475 was independently queried against three databases (prok_complete_genomes, refseq_genomes, and nt) using NCBI’s tBLASTn webserver and hits were downloaded in xml format. Using a Python script utilizing Biopython v. 1.7.8 functions (Cock et al., 2009), xml files were parsed and combined into a unique set of hits. Fully annotated complete genomes containing the homologs from the combined list of hits were downloaded from NCBI in genbank (gbk) format (~3GB).

Each genome was parsed and coding sequence (CDS) features of corresponding BLAST hits were extracted. Proteins with unique accession numbers and unique sequences (excluding pseudogenes) were retained and combined into a fasta file. Hits shorter than 200 amino acids were eliminated from the list to obtain the final set of homologs in distinct genetic backgrounds (Table 1).

### Phylogenetic tree generation

An iterative process was used to select the final set of 100 homologs and build the tree. The fasta file with all identified homologs was first submitted to the MAFFT webserver (used May 8^th^, 2021, https://mafft.cbrc.jp/alignment/server/) to obtain a multiple sequence alignment (MSA) with the E-INS-I option selected along with the remainder parameters set to their default values. Upon inspection of the MSA, low-scoring homologs that were also observed to be from adequately represented genera/species but introducing large gaps in the alignment were removed from consideration. The MSA computation was then repeated as above. Using this second MSA, a preliminary phylogenetic tree was generated with IQ-TREE21 multi-core v. 1.6.12 (Nguyen et al., 2014) with the optimal substitution model LG+R6 that provided the lowest Bayesian information criterion. One thousand ultra-fast bootstraps were performed to evaluate node support (options –bb 1000 –wbt). Upon inspection of the resulting tree, hard polytomies (Sayyari and Mirarab, 2018) that resulted in many extremely short branches were identified and, to preserve the phylogenetic diversity, only one representative homolog for each were retained to perform the final MAFFT alignment. A final MSA was computed with the remaining homologs (n=100 including SERP2475) and the final phylogenetic tree was computed using IQ-TREE with the same parameters noted above.

RAxML-NG v. 1.0.2 (Kozlov et al., 2019) was used to confirm the tree with similar corresponding parameters (100 bootstraps). Fig Tree v. 1.4.4 (http://tree.bio.ed.ac.uk/software/figtree/) was used for tree visualization, and Adobe Illustrator was used to overlay highlights and markings relevant to the study.

### Homolog neighborhood analysis

For the neighborhood analysis, the genomes containing the original 303 homologs (including SERP2475) were parsed and the amino acid sequences of neighboring proteins encoded on either side of the Blast hit (window sizes of 10, 20, and 30) were extracted into individual fasta files using a Python script. Pfam 34.0 database was downloaded (Pfam-A.hmm) and hmmpress (hmmer 3.3.2, hmmer.org) was used to index it. For each of the neighborhood fasta files, a Python script utilizing hmmscan was used to obtain and generate a new set of files that included protein family (pfam) predictions of its contents.

Predicted pfams were then searched against a set of 306 pfams with known defense-related functions (Table S3)—this list was compiled from the old and newly-identified defense pfams cited in (Gao et al., 2020). A Python script was then developed and utilized to determine the defense related neighbors for each homolog and analyzed to generate the plots for each neighbor window size.

### Phage adsorption and cell viability assays

Overnight cultures of *S. epidermidis* LM1680/p*SERP-NC* and LM1680/p*SERP-247*5 were diluted 1:100 in fresh BHI supplemented with antibiotics and 5 mM CaCl_2_, and incubated at 37°C with agitation for one hour. For the adsorption assay, Andhra or JBug18 were added to cultures (0.01:1 phage:cell ratio) and incubated at 37°C for 10 min. Cells along with adsorbed phages were pelleted at 8000 *x g* for 5 min at 4°C and resulting supernatants were passed through 0.45 mm syringe filter. The number of free phages in the supernatants were enumerated by the double-agar overlay method (Cater et al., 2017). The number of adsorbed phages were determined by subtracting the number of phages in suspension from the number that was initially added. Triplicate samples were prepared for each treatment, and two independent trials were conducted. For cell viability assays, 200 µl of the bacterial cultures were distributed into a 96-well microtiter plate (into triplicate wells for each treatment), and phages were added to cells at ratios of 1:1, 5:1, or 10:1. These bacteria-phage mixtures were incubated at 37°C with agitation for an additional five hours. 25 µl of 0.1% (w/v) 2,3,5 triphenyltetrazolium chloride (TTC) was added into each well and the microtiter plate was incubated at 37°C for an additional 30 mins to allow the colorless TTC to become enzymatically reduced to the red 1,3,5-triphenylformazan product by actively growing bacterial cells. The relative numbers of viable cells were then determined by measuring the absorbance at 540 nm. Triplicate measurements were averaged, and two independent trials were conducted.

### Phage infection time course followed by quantitative PCR

Phage infection time course assays were conducted in liquid media as previously described (Chou-Zheng and Hatoum-Aslan, 2019). Briefly, *S. epidermidis* LM1680 mid-log cells bearing p*SERP-NC* or *pSERP-2475* were infected with Andhra or JBug18 (phage:cell ratio of 0.5:1), cells were harvested at 0, 10, or 20 minutes post-infection, and their total DNA was extracted. Each qPCR reaction (25 µl) contained 500 ng of total DNA as template, 0.4 nM of phage-specific primers (N233 and N234) or host-specific primers (S001 and S002) (Table S7), and 1X PerfeCTa SYBR Green SuperMix (Quanta Biosciences). Separate standard reactions containing 10^2^–10^9^ DNA molecules were also prepared using purified Andhra phage DNA extract, JBug phage DNA extract, or bacterial genomic DNA extract. A CFX Connect Real-Time PCR Detection System (Bio-Rad) was used to amplify the DNA templates. Phage DNA copy number was normalized against host values, and the normalized value for the 0 min time point was set to one to obtain the relative DNA abundance for the rest of the time points as described previously (Chou-Zheng and Hatoum-Aslan, 2019). Triplicate measurements were taken for each of two independent trials.

### Construction of pET28b-10His-Smt3-based plasmids

pET28b-10His-Smt3-*SERP2475* and pET28b-10His-Smt3-*SERP2475-*230AA were constructed via Gibson assembly. Briefly, inserts were obtained by amplifying the complete and partial portion of *SERP2475* from *S. epidermidis* RP62a genome, and the backbone was amplified from a pET28b-His10Smt3 template.

Corresponding primers are listed in Table S7. The backbone PCR products were subjected to DpnI (NEB) digestion. Then, all PCR products were purified using the EZNA Cycle Pure Kit and Gibson assembled. Assembled constructs were introduced into *E. coli* DH5α via chemical transformation, and transformants were confirmed to have the desired plasmids using PCR and DNA sequencing with primers T7P and T7T (Table S7). Confirmed plasmids were purified using the EZNA Plasmid Mini Kit and introduced into *E. coli* Rosetta2 (DE3) cells for protein purification.

### Purification of recombinant SERP2475 and SERP2475-230AA

Recombinant proteins encoded in pET28b-10His-Smt3-based plasmids were overexpressed and purified as described previously with some modifications (Chou-Zheng and Hatoum-Aslan, 2019). Briefly, overnight cultures were diluted 1:100 in 1 L (for the truncated version) or 2 L (for the full-length version) of TB supplemented with appropriate antibiotics. Once the OD_600_ reached 0.5–0.6, cell-growth was arrested on ice for 20 minutes, and protein expression was induced with 0.3 mM isopropyl-1-thio-β-d-galactopyranoside (IPTG) and 2% ethanol. Induction proceeded 16-18 hr at 17°C with constant shaking. Cells were harvested and washed with cold PBS Buffer (137 mM NaCl, 2.7 mM KCl, 10 mM Na_2_HPO4, 1.8 mM KH_2_PO4). All subsequent steps were performed at 4°C. Each one-liter pellet was suspended in 30 ml of Buffer A (50 mM Tris–HCl, pH 9.0, 1.25 M NaCl, 200 mM Li_2_SO4, 10% sucrose, 15 mM Imidazole) containing one complete EDTA-free protease inhibitor tablet (Roche), 0.1 mg/ml lysozyme, and 0.1% Triton X-100. After 1 hr rotation, lysed cells were sonicated, and the insoluble materials were removed via centrifugation and filtration. Then, 3 ml (for 1-L pellet of truncated version), or 4 ml (for 2-L pellet of full-length version), of slurry Ni^2+^-NTA-agarose resin (ThermoFisher) was equilibrated with Buffer A and mixed with the cleared lysates. After 1 hr incubation, the resin was collected via centrifugation and washed with 40 ml of Buffer A. The resins containing the full-length protein from individual 1-L pellets were pooled. Resins were then transferred to a 5-ml gravity column (G-Biosciences) and further washed with 25 ml of Buffer A. Proteins were collected into 1-ml aliquots and eluted in stepwise elution with 3 ml of IMAC buffer (50 mM Tris–HCl pH 9.0, 250 mM NaCl, 10% glycerol) containing 50-, 100-, 200-, and 500-mM imidazole, respectively. The 10His-Smt3-tag was removed from most concentrated eluates using SUMO Protease (McLab, 1000 U) and supplied buffer (salt-free). Samples were dialyzed against IMAC buffer containing 25 mM Imidazole for 3 hr. Then, 2 ml (for the truncated version), or 1.5 ml (for the full-length version), of slurry Ni^2+^-NTA-agarose resin was equilibrated with IMAC Buffer containing 25mM Imidazole, and incubated with the dialysate for 1 hr. The resin was collected in a 5-ml gravity column, and tag-free proteins were collected and concentrated using a 10K MWCO centrifugal filter (PALL). Concentrated proteins were further purified by size exclusion chromatography using Superdex 75 Increase 10/300 GL (Cytiva) (for the truncated version) and Superdex 200 Increase 10/300 GL (Cytiva) (for the full-length version). Collected protein fractions were subjected to nuclease assays. Fractions with the highest peak concentration were combined and concentrated for nuclease time course and helicase assays. Proteins were resolved on 12% SDS-PAGE run at 120 V for 1.5 hours and visualized with Coomassie G-250, and the concentrations were determined using Bradford reagent (Bio-Rad).

### Nuclease assays

For exonuclease assays, single stranded DNA substrates were labeled on their 5’-ends by incubating with T4 polynucleotide kinase and γ-[^32^P]-ATP and purified over a G25 column (IBI Scientific). Radiolabeled substrates were combined with 7.5 µl of each protein fraction, or 130 pmols for time course assays, in 10 µl reactions containing nuclease buffer (25 mM Tris-HCl pH 7.5, 2 mM DTT) and 10 mM of MgCl_2_. Nuclease reactions were incubated at 37°C for 20 minutes (for experiments with different fractions), or 5, 10, and 20 minutes (for time course assays). Reactions were stopped by adding an equal volume of 95% formamide loading buffer and resolved on a 15% Urea PAGE gel at 55 W for 1.5 hours. Gels were exposed to a storage phosphor screen and visualized using an Amersham Typhoon biomolecular imager. For nickase assays, 250 ng of plasmids pBR322 or pUC19 (NEB) were combined with 15 µl of protein fractions, or 260 pmols for time course assays, in 20 µl reactions containing nuclease buffer (25 mM Tris-HCl pH 7.5, 2 mM DTT) and 10 mM of MgCl_2_. Nickase reactions were incubated at 37°C for 60 minutes (for experiments with different fractions), or 30, 60, 120, and 240 minutes (for time course assays). Reactions were stopped by placing on ice for 5 min, followed by incubation with 10 µg of proteinase K for 20 minutes at room temperature. Samples were then resolved on a 1% agarose gel run at 120 V for 50 minutes and visualized with ethidium bromide under UV transillumination with an Azure 400 imager.

### Helicase Assay

Double-stranded DNA duplexes were prepared by combining 5′-radiolabeled ssDNA oligonucleotides and unlabeled complementary ssDNA oligonucleotides (as shown in Fig. 5J) in a 1:2.5 molar ratio. The mixtures were heated to 95°C for 5 min and then slowly cooled down to room temperature over a period of 3 hours. The helicase assay was performed by first mixing radiolabeled DNA duplex with a 10-fold molar excess of unlabeled top-strand DNA to trap the complementary strand once unwound. This mixture was combined with 200 pmol SERP2475 in helicase buffer (25 mM Tris-HCl pH 9.0, 2 mM DTT, and 0.1 mg/ml BSA) supplemented with 2 mM MgCl_2_ and 5 mM ATP in a 50 µl reaction. Reaction mixtures were incubated at 37°C for 1h. As a separate positive control for unwinding, DNA substrates were heated to 95°C for 10 min in the absence of the enzyme. Reaction was stopped by adding 5 µl of the stop solution (0.1% [wt/vol] bromophenol blue, 0.1% [wt/vol] xylene cyanol, 8% [vol/vol] glycerol, 0.4% [wt/vol] SDS, 50 mM EDTA). Samples were resolved on an 8% (v/v) non-denaturing polyacrylamide gel at 130 V for 4 hours at 4°C. The gel was dried under vacuum at 80°C, exposed to a storage phosphor screen and visualized using an Amersham Typhoon biomolecular imager. Four independent trials were performed. ImageQuant software was used for densitometric analysis. The fraction of unwound (ssDNA) was determined using the following equation: density of ssDNA signal divided by the sum of densities of dsDNA and ssDNA signals. ***Mass spectrometry analysis*.** Protein bands corresponding to the 72 kDa (full-length) and 23 kDa (truncated) variants of SERP2475 in the FT fraction from the second step of purification were excised from a 12% SDS-PAGE gel stained with Coomassie G-250. Mass spectrometry was performed by the Cancer Center Mass Spectrometry and Proteomics Shared Facility at the University of Alabama, Birmingham. The bands were digested overnight with Trypsin Gold, Mass Spectrometry Grade (Promega, cat. #V5280) following the manufacturer’s instructions.

Peptide extracts were reconstituted in 0.1% formic acid/ddH_2_O at 0.1 μg/μl. Electrospray ionization tandem mass spectrometry was carried out, and the data were processed, filtered, grouped, and quantified, as previously reported in detail (Ludwig et al., 2016). The data were searched against a tailored database comprising of the *E. coli* proteome plus the protein sequence of interest (SERP2475).

### Phage hybrid generation and sequencing

JBug18-Andhra Hybrids 1-8 were isolated as immune resistant mutants following challenge of LM1680/p*SERP-2475* with a high titer lysate of JBug18 (~1 x 10^10^ pfu/ml). To generate JBug18-Andhra Hybrids 9-18, overnight cultures of *S. epidermidis* LM1680 harboring pT181*-gp03* or pT181*-gp0304* were diluted 1:100 in fresh TSB supplemented with antibiotics and 5 mM CaCl_2_. The mixture was incubated at 37°C for an hour with agitation, then JBug18 was added to the cells in a 1:1 ratio, and the incubation continued with agitation overnight. The next day, cells were pelleted by centrifugation at 8000 *x g* for 5 min and supernatant was filtered through 0.45 mm filter. Filtered lysates were mixed with LM1680-p*SERP2475* overnight culture (1:1) and the mixture was plated on TSA containing 5 mM CaCl_2_ using the double-agar overlay method (Cater et al., 2017). For all phage hybrids, individual plaques were isolated and re-plated three times on LM1680/p*SERP-2475* to purify. Phages were propagated and their DNA was extracted as previously described (Bari et al., 2017). Phage genomes were PCR amplified across the entire coding region for Hybrids 1-8 or *gp03-gp04* for Hybrids 9-18, and the PCR products were sequenced by the Sanger method (at Eurofins MWG Operon) using the primers listed in Table S7.

### Hybrid phage genome sequence analysis

For JBug18-Andhra Hybrids 1-8, Sanger sequencing reads covering their coding regions were manually assembled using SnapGene software. For JBug18-Andhra Hybrids 9-18, a single read covered the region of interest, therefore no assembly was required. Sequences for each set of hybrids (1-8 and 9-18) were aligned with corresponding genomic regions in Andhra and JBug18 using the Clustal Omega Multiple Sequence Alignment tool (https://www.ebi.ac.uk/Tools/msa/clustalo/). The sequence alignments (Supplementary files 2 and 3) were analyzed by a Python script developed in-house which first scans the alignment of JBug18 and Andhra, identifies each position of non-similarity, and then determines at those positions the fraction of hybrids that possess Andhra identity. The output data was exported into an Excel file, and the graphs showing the fraction of hybrids with Andhra identity at each position were generated using Microsoft Excel.

### Data availability

The raw phage sequence reads that support the findings of Figure 6 are publicly available through Figshare (<u>10.6084/m9.figshare.9598040</u>). The Accession codes for Andhra, JBug18, Pontiff, and Pike genomes are KY442063, MH972263, MH972262, and MH972261, respectively. Phages, mutant derivatives, and constructs can be made available upon written request.

### Code availability

The Python code library used for homolog and neighborhood analysis in Figure 2 is freely available at GitHub (https://github.com/ahatoum/Nhi). The Python code written to analyze the data for Figure 6 is freely available at GitHub (https://github.com/ahatoum/Hybrid-phage-genome-sequence-analysis).

## Supplementary Information

The following supplementary files are also available:

**Supplementary Figures 1-4**

**Table S1.** Accompanies figure 2. SERP2475 and 302 homologs in unique genetic backgrounds

**Table S2.** Accompanies figure 2. Protein families (Pfams) associated with anti-phage defense

**Table S3.** Accompanies figure 2. Defense-related Pfam neighbors within 20 genes of SERP2475 homologs

**Table S4.** Accompanies figure 2. Rank ordering of defense-related Pfams within 20 genes of SERP2475 and its homologs

**Table S5.** Accompanies figure 5C. Mass spectrometry analysis of proteins contained within “band a”.

**Table S6.** Accompanies figure 5C. Mass spectrometry analysis of proteins contained within “band b”.

**Table S7.** Oligonucleotides used for cloning and PCR in this study.

**Supplementary file 1.** SERP2475 and 99 selected homologs

**Supplementary file 2.** Multiple sequence alignment for JBug18-Andhra Hybrids 1-8

**Supplementary file 3.** Multiple sequence alignment for JBug18-Andhra Hybrids 9-18

## Acknowledgements

AH-A would like to acknowledge funding for this project from the National Science Foundation CAREER award [MCB/2054755] and an Investigators in the Pathogenesis of Infectious Disease Award from the Burroughs Wellcome Fund.

## Author Contributions

S.M.N.B., L.C-Z., O.H., K.C., V.D., and A.T., designed, created, and tested plasmid constructs; S.M.N.B. performed adsorption, cell viability, and phage hybridization assays; LCZ performed qPCR assays and analyses; S.M.N.B. and L.C-Z. performed protein purification; S.M.N.B., L.C-Z., and A.H-A. performed biochemical assays and analyses; B.A. performed all bioinformatic analyses; and A.H-A. conceived the study and wrote the manuscript. All authors have read and approved the manuscript.

## Supplementary Figures

**Figure S1.**
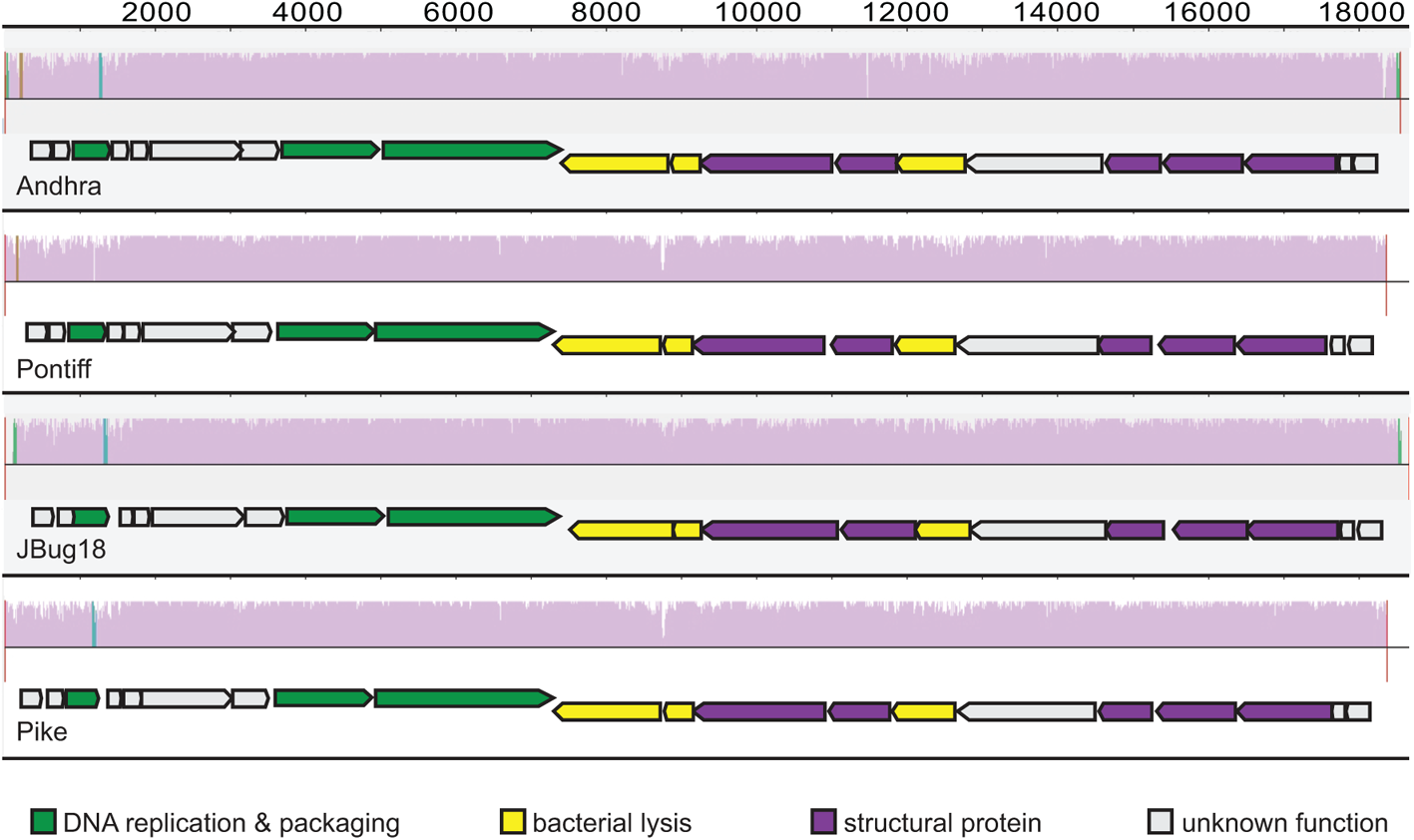
To accompany Figure 1. Four related *Podoviridae* phages with different host ranges. Shown is a multiple genome alignment of *S. epidermidis* podophages Andhra, Pontiff, JBug18, and Pike. Genome coordinates are shown on top, and colored histograms indicate the nucleotide similarity at each position derived from a multiple sequence alignment. The open reading frames for each phage are shown underneath the corresponding histogram. The histograms were generated using the MAUVE open source software (http://darlinglab.org/mauve/mauve.html) and the outlines of open reading frames from the MAUVE output were overlaid using Adobe Illustrator.

**Figure S2.**
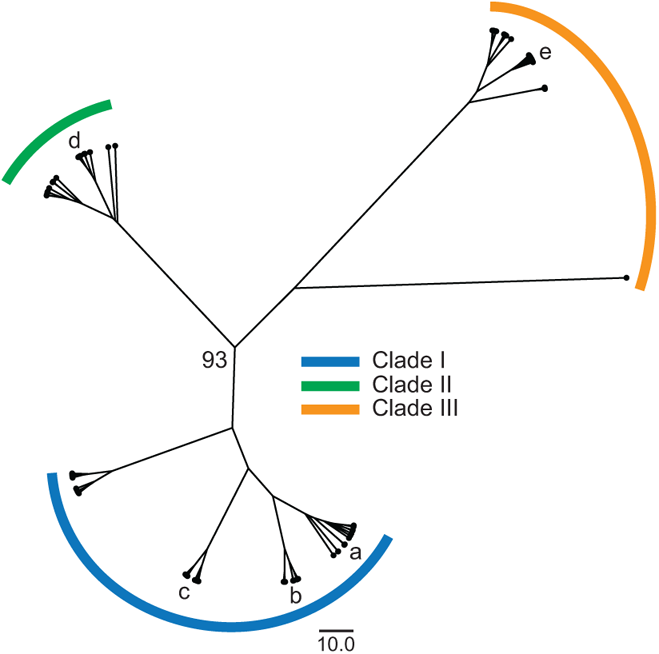
To accompany Figure 2. A dendrogram generated for SERP2475 and 99 homologs. Positions of homologs selected for functional characterization are indicated with lower-case letters: a. WP_045177897 from *S. aureus* MJ163; b. WP_002489608 (i.e. SERP2475) from *S. epidermidis* RP62a; c. WP_000632676 from *S. aureus* CA-347; d. WP_013870910 from *Lacinutrix* 5H-3-7-4; e. WP_101958732 from *V. vulnificus* FORC54.

**Figure S3.**
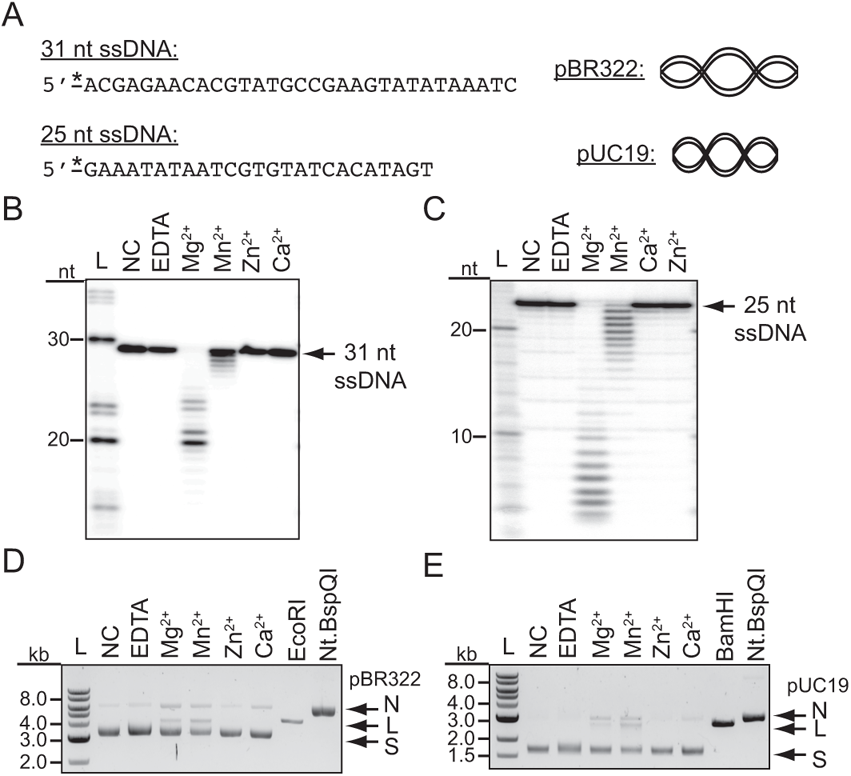
To accompany Figure 5. Activity of SERP2475 N-terminal 230 amino acids in the presence of different metals and substrates. (**A**) Illustration of different substrates used for nuclease assays. Asterisks indicate the position of the radiolabel on the 5’-end of linear substrates. Exonuclease assays (**B** and **C**) and nickase assays (**D** and **E**) are shown in which various substrates were combined with a purified preparation of the N-terminal 230 AAs of SERP2475 (13 nM) in a reaction supplemented with indicated metals or EDTA (2 mM) and incubated at 37°C for 20 min (for exonuclease assays) or 1 h (for nickase assays). Shown are representative images for three independent trials.

**Figure S4.**
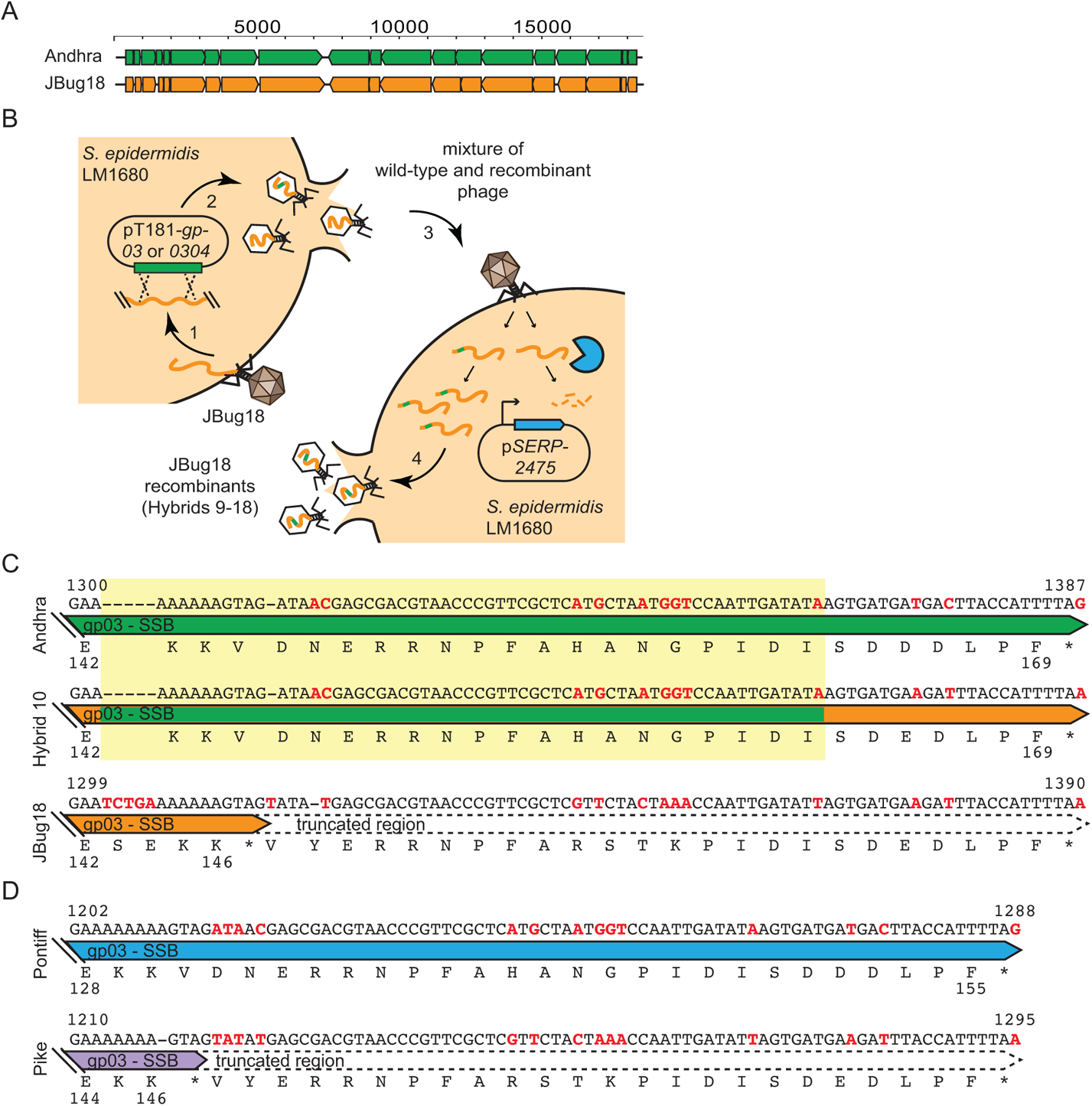
Accompanies Fig. 6. Generation of Hybrids 9-18 and comparison of SSB sequences. (**A**) A pairwise comparison of the open reading frames of Andhra and JBug18. (**B**) A diagram of the method used to generate JBug18-Andhra Hybrids 9-18. (**C**) Sequence comparison between the SSBs of Andhra, JBug18, and Hybrid 10, which gained resistance to immunity through the acquisition of only 60 nucleotides of Andhra-derived sequence (highlighted in yellow). (**D**) a similar comparison between the SSBs of phages Pontiff and Pike.

